# Differential patterns of gyral and sulcal morphological changes during normal aging process

**DOI:** 10.1101/2020.10.30.361626

**Authors:** Hsin-Yu Lin, Chu-Chung Huang, Kun-Hsien Chou, Albert C. Yang, Chun-Yi Zac Lo, Shih-Jen Tsai, Ching-Po Lin

## Abstract

The cerebral cortex is a highly convoluted structure with distinct morphologic features, namely the gyri and sulci, which are associated with the functional segregation or integration in the human brain. During the lifespan, the brain atrophy that is accompanied by cognitive decline is a well-accepted aging phenotype. However, the detailed patterns of cortical folding change during aging, especially the changing trajectories of gyri and sulci, which is essential to brain functioning, remain unclear. In this study, we investigated the morphology of the gyral and sulcal regions from pial and white matter surfaces using MR imaging data of 417 healthy participants across the lifespan (21-92y). To elucidate the age-related changes in the cortical pattern, we fitted cortical thickness and intrinsic curvature of gyri and sulci using the quadratic model to evaluate their trajectories during normal aging. Our findings show that comparing to gyri, the sulcal thinning is the most prominent pattern during the aging process, and the gyrification of pial and white matter surfaces were also affected differently, which implies the vulnerability of functional segregation during aging. Taken together, we propose a morphological model of aging that may provide a framework for understanding the mechanisms underlying the gray matter degeneration.

## Introduction

The morphology and function of the human brain change throughout the aging process. However, the mechanisms underlying how the structures of gyri and sulci are altered with age remain unclear. In recent decades, in vivo magnetic resonance imaging (MRI) has been widely utilized to investigate the effects of aging on the human brain (Good et al., 2001; Jernigan et al., 2001; Raz et al., 2004; Salat et al., 2004), and these studies have provided information on how the structures of the human brain change during the course of a lifespan. The decrease in cortical volume is the most dramatic change that occurs during aging (Scahill et al., 2003). Nevertheless, the highly convoluted and complex nature of the cerebral cortex implies aspects such as folding and thickness of the cortex may have different influences on cortical morphology and brain function. In that case, in addition to volumetric measurements, surface area, gyrification, and thickness measurements also provide detailed information for brain morphology (Gautam et al., 2015; Panizzon et al., 2009; Winkler et al., 2010). The cerebrum attains its folding structures through a complex orchestrated set of systematic mechanisms including mechanical tension, differential proliferation, mechanical buckling, and differential expansion (Van Essen et al., 2018). Previous studies have suggested that the development of cortical anatomy is dominated by genetic factors rather than random convolutions (Peng et al., 2016). These genetic factors could contribute to changes of cortical structure throughout the lifespan (Fjell et al., 2015; Ronan and Fletcher, 2015). As such, previous findings suggest that cortical features might also degenerate in a non-random, systematic way.

It is worth noting that the locations of specific gyri and sulci (e.g. central sulcus, visual cortex…) are consistent in different individuals and correspond to regional functions, and these structures are considered structural-functional entities (Brodmann, 1909; Welker, 1990). With this point of view, gyrification and cortical thickness are thought to reflect the functional aspects of the cortex, such as intelligence, behavior complexity and cognition (Gautam et al., 2015; Kaas, 2013). Gyrification is thought to be a mechanical process that causes the surface to buckle (Ronan et al., 2014; Van Essen et al., 2018), although the primary underlying mechanism is still under debate (Xu et al., 2010). Traditionally, local gyrification index (LGI) (Schaer et al., 2008) has been used as a proxy to probe the regional folding degree of the human brain (Nanda et al. 2014, Zhang et al. 2014). However, due to the ratio-principle of LGI, it has less sensitivity compared to the intrinsic curvature on describing the complex pattern of cortical surface, especially in the deep brain regions, e.g. the insula (Griffin, 1994; Ronan et al., 2011). It has been suggested that intrinsic curvature is a more sensitive index to describe cortical folding and is presumed to reflect the differential development and cortical connectivity in different areas of the cortex (Ronan et al., 2014). Intrinsic curvature of the pial and white matter surface has also been found to be related to the structural changes of both superficial or deeper layers of cortex (Ronan et al., 2012; Wagstyl et al., 2016). Cortical thickness is a common feature reflecting total neuronal body in the cortex. Sulci, which are thinner than gyri, communicate locally with neighboring structures, while gyri act as functional centers that connect remote gyri to neighboring sulci (Deng et al., 2014; Fischl and Dale, 2000). Stronger functional connectivity was found in gyri than sulci, as supported by a series of experiments (Deng et al., 2014). Previous studies have shown that cognitive performance is associated with pattern changes in regional gyri (Gregory et al., 2016; Jones et al., 2006; Turner and Spreng, 2012), and that the variability of sulcal volume is found to be related to the progression of the neurological diseases in patients (Im et al., 2008; Mega et al., 1998; Sullivan et al., 1998). In the healthy neurodevelopmental process, cortex nonuniformly thins and demonstrates increased thinning in sulci compared to gyri (Vandekar et al., 2015). Moreover, gyri and sulci exhibit opposite trends in curvature changes during aging (Magnotta et al., 1999) and show different kinds of specialized organizations and connectivity (Welker, 1990). Together, the effects of aging on the regional thickness and gyrification of the brain result in decreased cognitive functions (Gregory et al., 2016; Salat et al., 2004). However, sulci are recently deemed to be more functionally segregated in structural connectivity and the increasing age is accompanied by decreasing segregation in large-scale brain systems (Chan et al., 2014; Liu et al., 2017). The morphological evidence for the mechanism underlying such degeneration of the human brain reported by the studies mentioned above and others is not unified. Thus, investigations of distinguishing gyral and sulcal morphology and comparing them are critical.

Previous studies have suggested that the relationship between cortical degeneration and advancing age is not linear, and the degeneration trajectories of cortical features might also differ (Cao et al., 2017; Klein et al., 2014; Storsve et al., 2014). Therefore, understanding the variations in brain morphology and its trajectories throughout the lifetime is a crucial step towards revealing the epicenter of aging-related degeneration of cortical morphology, because the pattern may reflect the configuration of connectivity of the brain (Ronan et al., 2011). By using the image dataset that was collected by a single scanner with a large sample size and a broad age range, this study aimed to investigate the following three issues at both a whole-brain and a regional level: 1) to reassure whether the effects of age on the cortical morphological features are nonlinear across the lifespan, 2) whether age differentially and/or systematically affects gyri and sulci, 3) whether the pial and white matter surfaces display distinct variations in their patterns during aging. We believe that measuring detailed morphological features to depict the degenerative pattern of the cerebral cortex could help elucidate underlying degeneration mechanisms.

## Materials and Methods

### Participant characteristics and image data acquisition

A total of 417 healthy Chinese controls were recruited from northern Taiwan, and their ages ranged from 21 to 92 (Male/Female: 211/206). All of the included participants had sufficient visual and auditory acuity to undergo basic cognitive assessment. This research was conducted in accordance with the Declaration of Helsinki and approved by the Institutional Review Board of Taipei Veterans General Hospital. Written informed consent was obtained from all participants after they had been given an overview of this study. A trained research assistant used the diagnostic structured Mini-International Neuropsychiatric Interview (M.I.N.I.) to evaluate each subject (Sheehan et al., 1998). The participants’ cognitive functions were assessed using the Mini-Mental State Examination (MMSE) (Folstein et al., 1975), Wechsler Digit Span tasks (forward and backward), and Clinical Dementia Rating (CDR) scale (Hughes et al., 1982) to avoid enrolling participants with possible dementia or cognitive impairments. Subjects who met any of the following exclusion criteria were not enrolled in the study: (1) any Axis I psychiatric diagnoses on the Diagnostic and Statistical Manual of Mental Disorders-IV (American Psychiatric, 2000), such as mood disorders or psychotic disorders; (2) any neurological disorders, such as dementia, head injury, stroke, or Parkinson’s disease; (3) illiterate; (4) an MMSE score less than 24; and/or (5) elderly (age of 65 or over) with a CDR score over 0.5. The demographic information for the subjects is listed in Table 1.

**Table 1.**
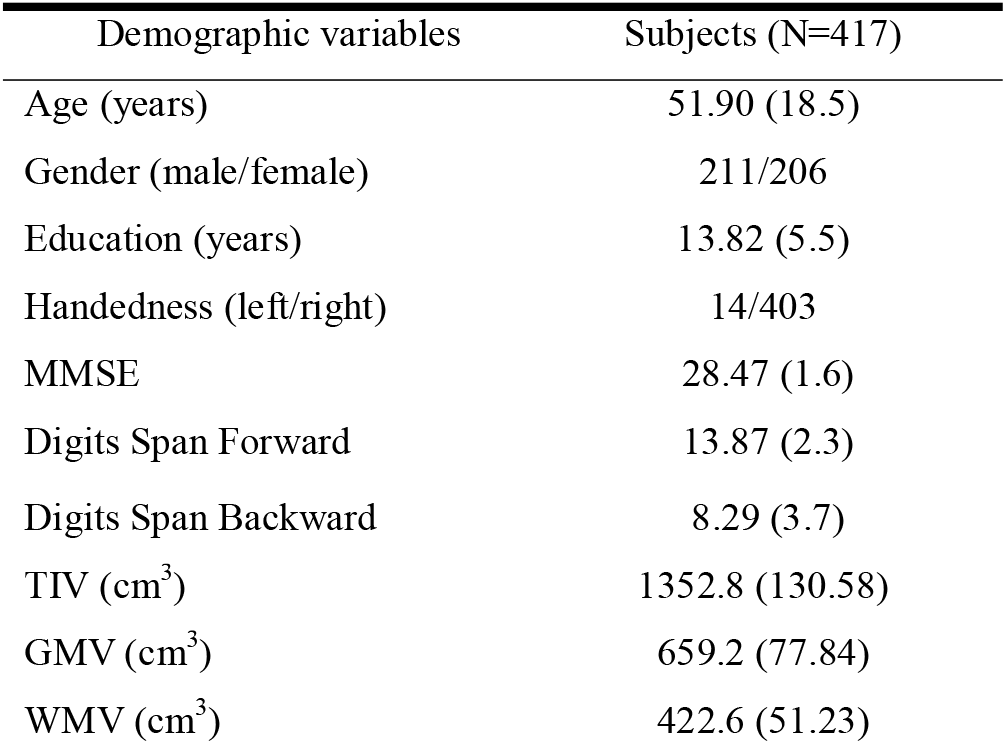
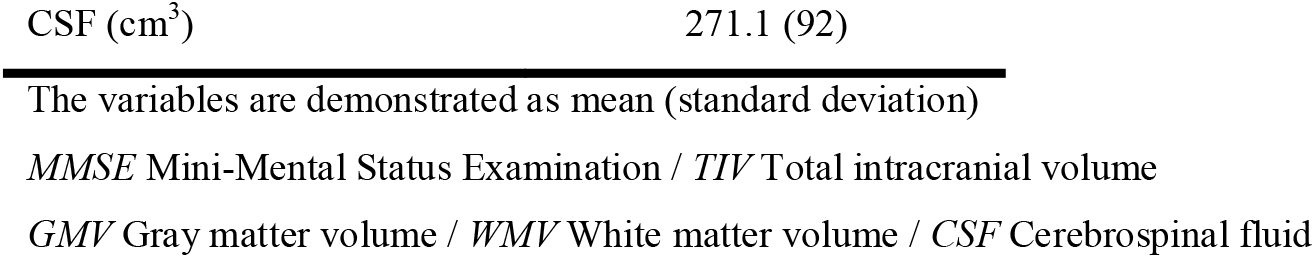
Demographic of included healthy participants

| Demographic variables | Subjects (N=417) |
| --- | --- |
| Age (years) | 51.90 (18.5) |
| Gender (male/female) | 211/206 |
| Education (years) | 13.82 (5.5) |
| Handedness (left/right) | 14/403 |
| MMSE | 28.47 (1.6) |
| Digits Span Forward | 13.87 (2.3) |
| Digits Span Backward | 8.29 (3.7) |
| TIV (cm<sup>3</sup>) | 1352.8 (130.58) |
| GMV (cm<sup>3</sup>) | 659.2 (77.84) |
| WMV (cm<sup>3</sup>) | 422.6 (51.23) |

The variables are demonstrated as mean (standard deviation)
*MMSE* Mini-Mental Status Examination / *TIV* Total intracranial volume
*GMV* Gray matter volume / *WMV* White matter volume / *CSF* Cerebrospinal fluid

All MRI scans were performed on a 3 Tesla Siemens scanner (MAGNETOM Trio TIM system, Siemens AG, Erlangen, Germany) with a 12-channel head coil at National Yang-Ming University. High-resolution structural T1-weighted (T1w) MRI scans were acquired with a three-dimensional magnetization-prepared rapid gradient echo sequence (repetition/echo time = 2,530/3.5, inversion time = 1,100 ms, field of view = 256 mm, flip angle =7°, matrix size = 256 × 256, 192 sagittal slices, isotropic voxel size= 1 mm^3^, and no gap). Each participant’s head was immobilized with cushions inside the coil to minimize the generation of motion artifacts during image acquisition.

### Cortical reconstruction

The pial and white matter surfaces of each subject were reconstructed using an automated cortical surface reconstruction approach in FreeSurfer 5.3 (http://surfer.nmr.mgh.harvard.edu) according to the following steps: registration, skull-stripping, segmentation of the gray matter, white matter and cerebrospinal fluid, tessellation of the gray-white matter boundary, automated topology correction, and surface deformation according to intensity gradients to optimally place the gray-white and gray-cerebrospinal fluid borders at the locations of the greatest shifts in intensity, which defines the transition to the other tissue class (Dale et al., 1999). The vertices were arranged in a triangular grid with the spacing of approximately 1 mm (approximately 16,000 grid points) in each hemisphere. For any inaccuracies, the qualities of the segmentation and surface reconstruction were checked carefully by four researchers using the double-blinded method (The dataset was divided into four parts. Each of the researchers checked through two parts of the datasets, and every part was checked twice by two different people.) The data of the subjects were excluded if two people marked it as poor quality. For those only marked once by the researchers, we checked the data again and decided whether it should be removed from further processing and analyses. A total of 15 subjects were excluded in the end because of the poor quality. We separated both the left and right hemispheres into gyral and sulcal regions using FreeSurfer-generated mean curvature (Sulci: mean curvature value > 0/Gyri: mean curvature value < 0) for further comparison.

### Cortical thickness

By calculating the shortest distance between the pial surface and the gray matter-white matter boundary of the tessellated surface, we obtained the vertex-wise cortical thickness. To observe an overall trend of thickness, we applied a vertex-wise analysis, and the surface was smoothed by a 15-mm Gaussian kernel. We further calculated the average thicknesses of the gyral and sulcal regions and the ratio of the gyral and sulcal thicknesses to investigate the differential effects of aging on these two structural features.

### Intrinsic curvature

Intrinsic curvature is a fundamental property of a surface, and its measurements reflect higher complexity intrinsic information of surfaces. The degree of intrinsic curvature is dependent on the degree of the differential, with a bigger differential resulting in a greater degree of curvature, which might reflect the underlying connectivity of the human cortex (Ronan et al., 2014). The higher the intrinsic curvature value, the higher the complexity, underlying connectivity, and folding or curves are in one region.

The vertex-wise intrinsic curvature was calculated using Caret software (v5.65, https://www.nitrc.org/projects/caret/) as the product of the principal curvatures (Ronan et al., 2011; Ronan et al., 2012). The post-processing and filtration of the curvature were done in MATLAB (The MathWorks, Inc., Natick, MA, USA). The modulus of intrinsic curvature was calculated at the pial and white matter surfaces. A low-pass filter (threshold = 2 mm^2^) was applied to minimize error, keep the curvature values compatible with the resolution of the cortical reconstruction and remove the abnormal values (Ronan et al., 2011). Additionally, the intrinsic curvature of the pial and white matter surfaces was separately extracted for the statistical analyses. To detect an overall trend, a vertex-wise analysis was applied, and the surface was smoothed by a 15-mm Gaussian kernel. Next, we calculated the average intrinsic curvatures for gyral and sulcal regions and the ratio of the gyri/sulci at the pial and white matter surfaces to investigate the differential aging effects.

### Regional analysis

We performed a regional analysis by dividing the cortical surface of each subject into five lobes: frontal, temporal, parietal, occipital, and cingulate. The lobes of the brain were defined by the Desikan-Killiany atlas (Desikan et al., 2006). The average gyri, sulci, and gyri/sulci ratio of thickness and intrinsic curvature were calculated in each lobe.

### Statistical analyses and curve fitting

For each hemisphere, an age correlation analysis of cortical thickness and intrinsic curvature (on the pial and white matter surfaces separately) was first tested using the General Linear Model vertex-by-vertex with gender and total intracranial volume as covariates in order to reveal any general variation trends on the brain surface.

The linear, quadratic and cubic models were applied in the regression analyses across age to determine the aging trajectory of cortical thickness (gyral thickness, sulcal thickness, and gyri/sulci thickness ratio) and intrinsic curvature (gyral intrinsic curvature, sulcal intrinsic curvature, and gyri/sulci intrinsic curvature ratio) on the pial and white matter surfaces separately. For model selection, the linear (y = p*age + p1), quadratic (y = p*age^2 + p1*age + p2), and cubic (y = p*age^3 + p1*age^2 + p2*age + p3) models were all tested using averaged whole brain measurements, and we chose the better model according to the properties of goodness-of-fit, including the Akaike information criterion (AIC) (Akaike, 1974), root mean square error (RMSE), and R^2^. To test the significance (*p* < .05) of each fitted model and minimize the type-I errors, we applied permutation-based multiple testing on all aging trajectories, reassigning age randomly 10,000 times.

### Relationships between cognitive performance and structural measures

Hierarchical multiple regression analysis was performed in SPSS to examine whether the general cognitive performance (MMSE/Digit span tasks, as dependent variable) can be explained when using the structural measures of gyri and sulci on the pial and white matter surfaces as additional independent variables, while age, gender and education were firstly entered in the regression model as covariates of non-interest. To test the significance of the regression models, we used a p-value threshold of 0.05.

### Stepwise regression predicting age using the structural measurements as predictors

Stepwise regression analysis was performed to determine the relative contribution of the structural measurement to chronological aging. Sex and TIV were forced to enter in the stepwise model, and the six structural measurements including gyral/sulcal thickness and intrinsic curvature in the pial or white surface were then entered to the stepwise model using a selection criterion of p < .05.

## Results

### Aging effect in brain tissue volume

To ensure that the recruited samples are in the aging process without muddling obvious development of brain tissues, we firstly examined the relationship between brain tissue volume (GMV, WMV, CSFV, and TIV) and age across 21-92 years old. The GMV, WMV, CSFV, and TIV are plotted as a function of age in supplementary Fig. S1. We confirmed that GMV and WMV decrease with age, CSFV increase with age and no correlation between TIV and age was observed in the analyzed sample.

### Vertex-wise linear correlations between age and cortical measurements

Across the age range of 21–92 years, vertex-wise cortical thickness showed a global negative linear correlation with age in both hemispheres (Fig. 1a). The vertex-wise age correlation results showed a regional decline (uncorrected *p* < 0.01) of the intrinsic curvature on the pial surface (Fig. 1b). However, the intrinsic curvature had an overall increasing pattern (uncorrected *p* < 0.01) on the white matter surface (Fig. 1c).

**Fig. 1.**
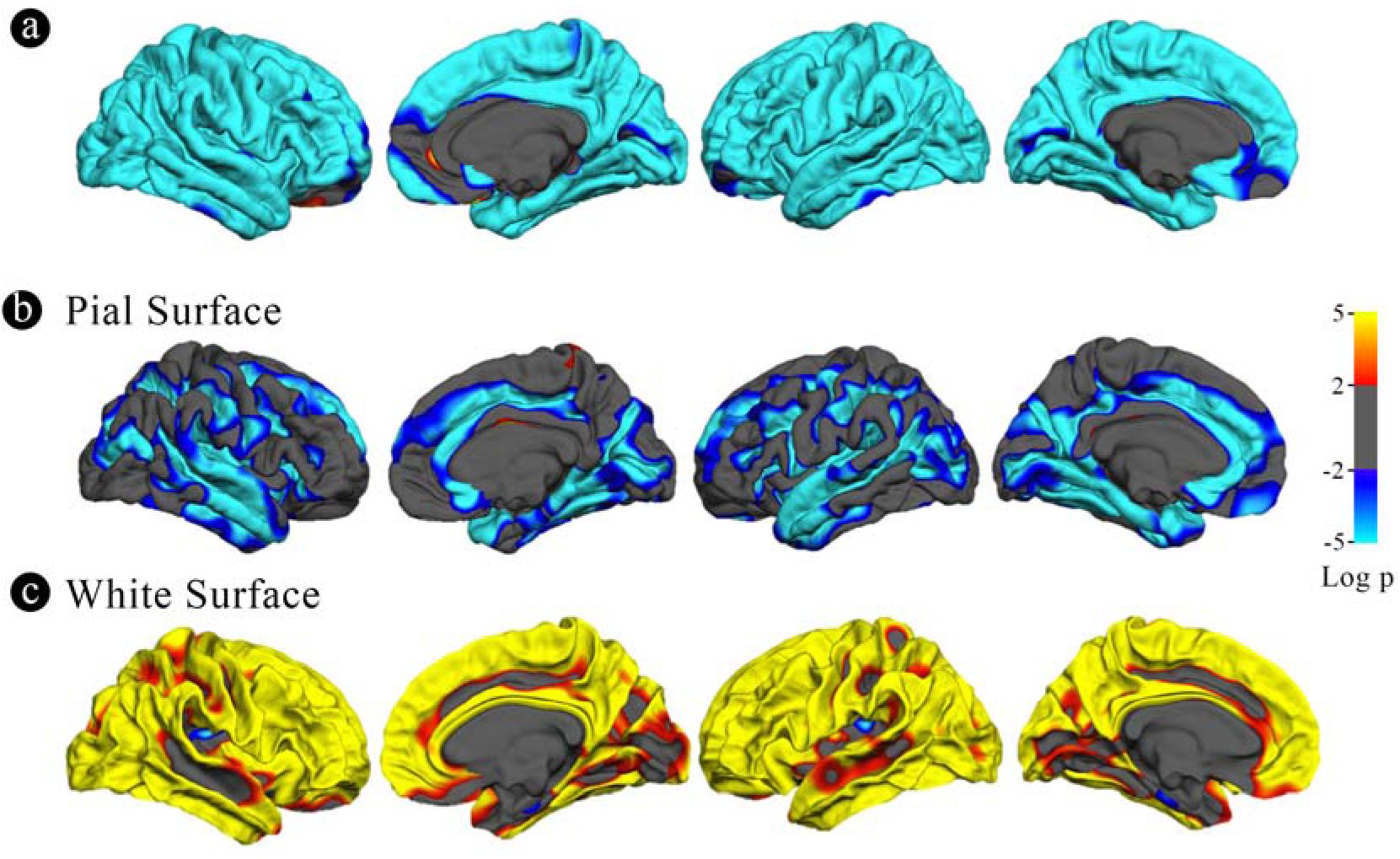
(a) Vertex-wise age correlation in cortical thickness. (b)(c) Vertex-wise age correlation in intrinsic curvature on the pial and white matter surfaces. The threshold was set to 2. Vertices with *p* < .01, uncorrected, are colored. Blue indicates a negative correlation; red indicates a positive correlation.

### Goodness-of-fit tests of the polynomial regression models

We examined the goodness-of-fit of the linear, quadratic, and cubic models, including looking at the two parameters: Root Mean Square Error (RMSE) and Akaike Information Criterion (AIC). For cortical thickness, quadratic models stood out to be the best fit (Supplementary Table S1). On the pial surface, bigger differences were found between the linear and quadratic models in both the RMSE and AIC results (Supplementary Table S2). The parameters of the quadratic and cubic models showed nearly the same, but the cubic values were smaller in most cases. The goodness-of-fit profile on the white matter surface was identical to that of the pial surface (Supplementary Table S3). The RMSE and AIC values were steady and smaller in the quadratic and cubic models compared with those of the linear model. Although some parameters were smaller using the cubic model compared with the quadratic model, those of cubic and quadratic models were very similar. Hence, we integrated the findings and only looked at the quadratic effects in the following analyses, which illustrated the aging process better but were not over-fit (For the linear results, see Supplementary Fig. S2 and Table S4).

### Quadratic relationship between age and whole-brain cortical measurements

All fitted curves were tested using permutation tests, and the results were all significant (*p* < 0.05). According to the quadratic regression results (Fig. 2, Table 2), both the cortical thicknesses of the gyri and sulci decreased with age [Gyral R^2^ (lh/rh): 0.400/0.370; Sulcal R^2^ (lh/rh): 0.562/0.526]. The gyri/sulci thickness ratio increased with age [R^2^ (lh/rh): 0.287/0.231], which implied that the degree of decrease in the sulci was larger than that of the gyri.

**Fig. 2.**
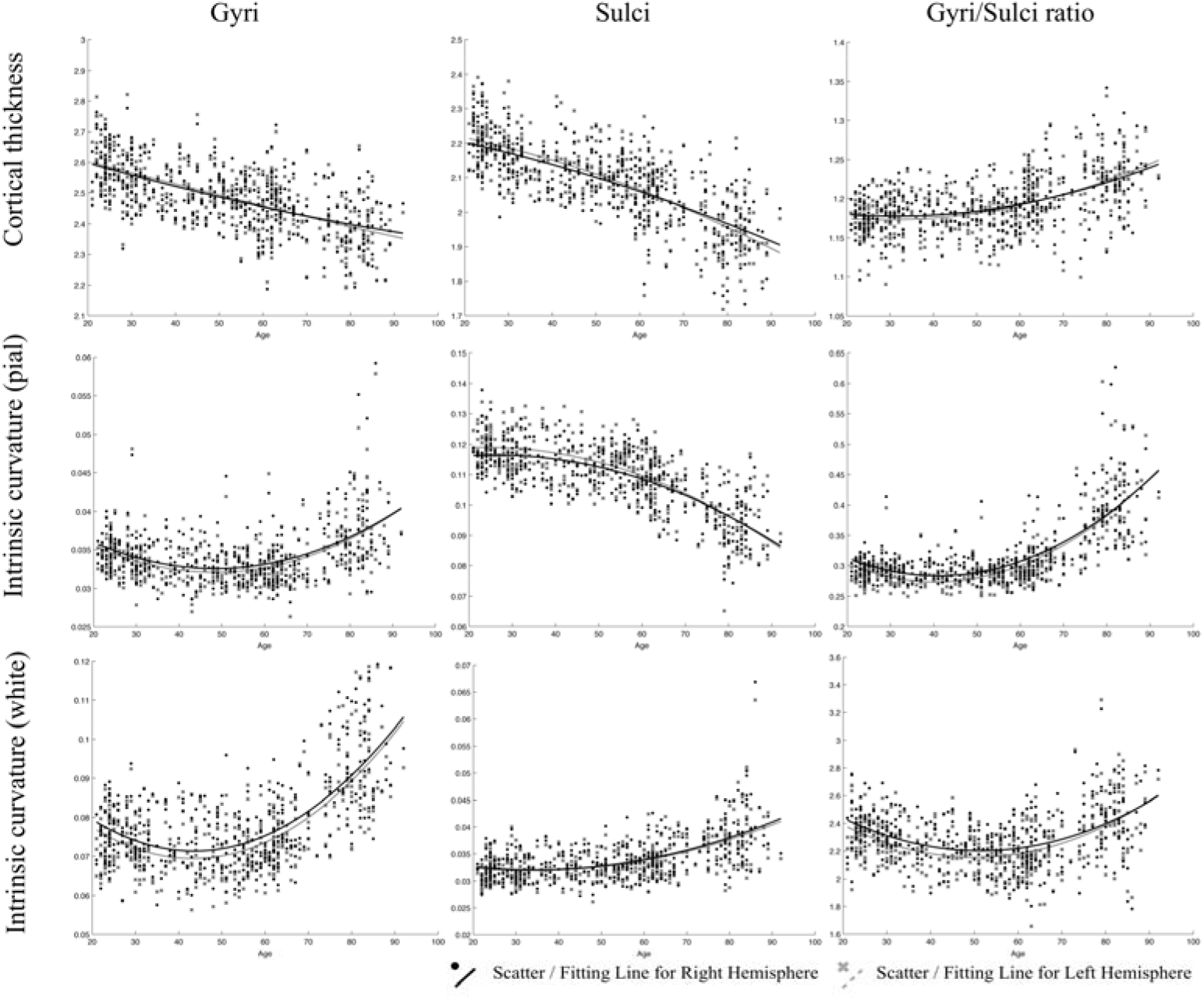
Results of the regressions of age with gyri, sulci, and the gyri/sulci ratio of cortical thickness and intrinsic curvature on the pial and white matter surfaces. The lines refer to the fitted curve for the age and measurements, and the dots indicate the distribution of that data for the subjects.

**Table 2.**
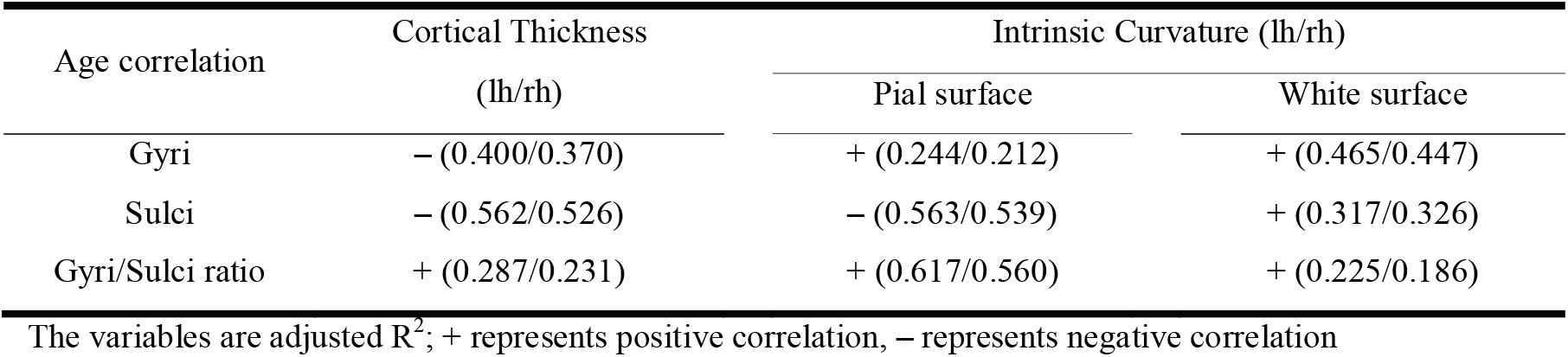
Results of quadratic regression analyses

| Age correlation | Cortical Thickness | Intrinsic Curvature (lh/rh) |  |
| --- | --- | --- | --- |
|  | (lh/rh) | Pial surface | White surface |
| Gyri | – (0.400/0.370) | + (0.244/0.212) | + (0.465/0.447) |
| Sulci | – (0.562/0.526) | – (0.563/0.539) | + (0.317/0.326) |
| Gyri/Sulci ratio | + (0.287/0.231) | + (0.617/0.560) | + (0.225/0.186) |

The variables are adjusted $R^2$ ; + represents positive correlation, – represents negative correlation

For gyrification, the aging process affected the gyri and sulci in opposite ways on the pial surface: a negative quadratic correlation was found with age in the sulcal region [R^2^ (lh/rh): 0.563/0.539], while a positive quadratic correlation was found with age in the gyral region [R^2^ (lh/rh): 0.244/0.212]. Both the gyral and sulcal intrinsic curvatures of the white matter surface increased quadratically with age [Gyral R^2^ (lh/rh): 0.465/0.447; Sulcal R^2^ (lh/rh): 0.317/0.326]. The gyri/sulci intrinsic curvature ratio on the white matter surface increased with age [R^2^ (lh/rh): 0.225/0.186], which indicated that the degree of increase of the sulci was smaller than that of the gyri. The results were all similar and consistent between the right and left hemispheres. The average thickness and intrinsic curvature values are shown in Supplementary Table S5.

### Quadratic relationship between age and cortical measurements of the five lobes

The relationship between cortical measurements and age for the five lobes including frontal, parietal, temporal, occipital lobe, and cingulate were examined. Most of the age-cortical measure relationships in the lobes were consistent with the whole-brain results, except for the gyri/sulci ratio of thickness in the right frontal lobe and left cingulate. The quadratic relationships between age and the sulcal intrinsic curvature on the white matter surface of the occipital lobe were not significant (*p* > 0.05) (Table 3, Fig. S3–S7).

**Table 3.** Results of quadratic regression analyses of the 5 lobes

| Lobe | Age correlation | Cortical Thickness<br>(lh/rh) | Intrinsic Curvature (lh/rh) |  |
| --- | --- | --- | --- | --- |
|  |  |  | Pial surface | White surface |
| Frontal | Gyri | − (0.414/0.259) | + (0.245/0.271) | + (0.448/0.457) |
|  | Sulci | − (0.284/0.271) | − (0.366/0.315) | + (0.387/0.391) |
|  | Gyri/Sulci ratio | + (0.078/ N.A.) | + (0.494/0.485) | + (0.100/0.098) |
| Parietal | Gyri | − (0.414/0.259) | + (0.192/0.185) | + (0.461/0.446) |
|  | Sulci | − (0.483/0.478) | − (0.443/0.436) | + (0.223/0.201) |
|  | Gyri/Sulci ratio | + (0.261/0.248) | + (0.561/0.545) | + (0.243/0.187) |
| Temporal | Gyri | − (0.350/0.353) | + (0.228/0.197) | + (0.408/0.388) |
|  | Sulci | − (0.623/0.594) | − (0.609/0.561) | + (0.220/0.198) |
|  | Gyri/Sulci ratio | + (0.493/0.426) | + (0.585/0.498) | + (0.202/0.157) |
| Occipital | Gyri | − (0.117/0.142) | + (0.096/0.030) | + (0.324/0.251) |
|  | Sulci | − (0.561/0.583) | − (0.386/0.358) | N.A. |
|  | Gyri/Sulci ratio | + (0.513/0.490) | + (0.418/0.314) | + (0.299/0.263) |
| Cingulate | Gyri | − (0.414/0.259) | + (0.137/0.091) | + (0.358/0.220) |
|  | Sulci | − (0.284/0.271) | − (0.378/0.368) | + (0.143/0.295) |
|  | Gyri/Sulci ratio | + (0.005/ N.A.) | + (0.413/0.218) | + (0.088/0.121) |

The significance of the curve fitting is evaluated using the permutation test, only the $R^2$ for the significant curves were shown in the table ( $p < .05$ ). The variables are adjusted $R^2$ ; + represents positive correlation, − represents negative correlation.

### Relationships between cognitive performance and structural measures

The hierarchical regression models were found significant between MMSE and sulcal cortical thickness and gyral intrinsic curvature on the white matter surface. In the first step, the regression model including age, gender and education was significant (R^2^= .236, p < .001). In the second step, adding the whole brain sulcal cortical thickness into the model explained additional 8% of variance, and the model stayed significant (R^2^ = .244, F-change p = .041). In the third step, adding the whole brain gyral intrinsic curvature into the model also explained additional 8% of variance, and the model was significant (R^2^ = .252, F-change p = .039). The models at the second and third steps revealed that MMSE was positively correlated with the cortical measurements (Fig. 3; sulcal cortical thickness: β = .141, p = .029; gyral intrinsic curvature: β = .109, p = .039).

**Fig. 3.**
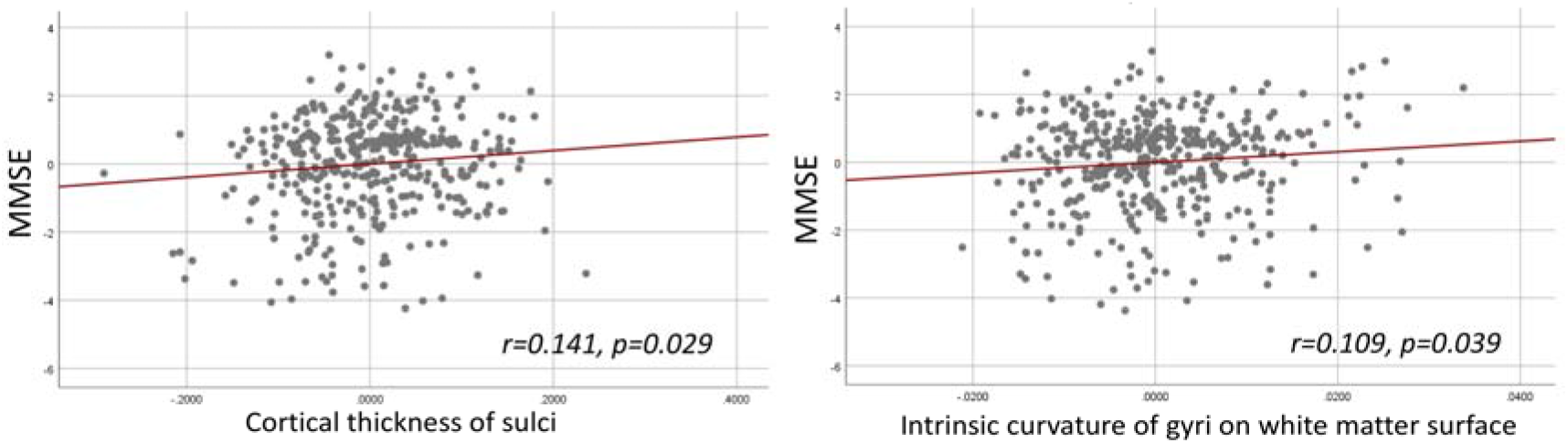
Hierarchical multiple regression result. Left: Partial regression plot of MMSE and sulcal cortical thickness. Right: Partial regression plot of MMSE and gyral intrinsic curvature of the white matter surface.

### Different contribution of the structural measurements in chronological aging trajectory

By using stepwise regression that adjusted for the sex and TIV effect, four of the six measurements were selected into the final model with adjusted, including mean thickness of sulci (standardized β = −.214, p < .001), mean intrinsic curvature of the pial (standardized β = −.524, p < .001) and white surface of sulci (standardized β = .544, p < .001), and mean intrinsic curvature of pial surface of gyri (standardized β = −.284, p < .001).

### The aging model

To summarize the various measurements and structural findings in the current and previous studies, we suggest a putative model of the general pattern of cortical aging (Fig. 4). In this model, the gyral intrinsic curvature of the pial surface increases slightly with age, while the tips of the gyri stay close to the skull (Fig. 4-1). The gyral intrinsic curvature on the white matter surface increases with age so that the surface moves outward, resulting in a decrease in cortical thickness (Fig. 4-2). The sulcal intrinsic curvature of the pial surface decreases with age (Fig. 4-3), which indicates that the sulci are getting wider and flatter on the surface with increasing age. The pial border also moves outward, resulting in a decline in the sulcal depth and increase in sulcal width. On the white matter surface, the sulcal intrinsic curvature increases with age, and the gray-white matter boundary moves outward and becomes cusped (Fig. 4-4). The gyral intrinsic curvature on the white matter surface increases more intensely with age than the sulcal curvature does. Finally, the decreasing extent of the sulcal thickness appears larger than that of the gyral thickness. Additionally, we found that the trends of all trajectories were consistent for both gender (see Supplementary Fig. S8), indicating that the overall pattern of the current putative model reflects common mechanisms of change during aging.

**Fig. 4.**
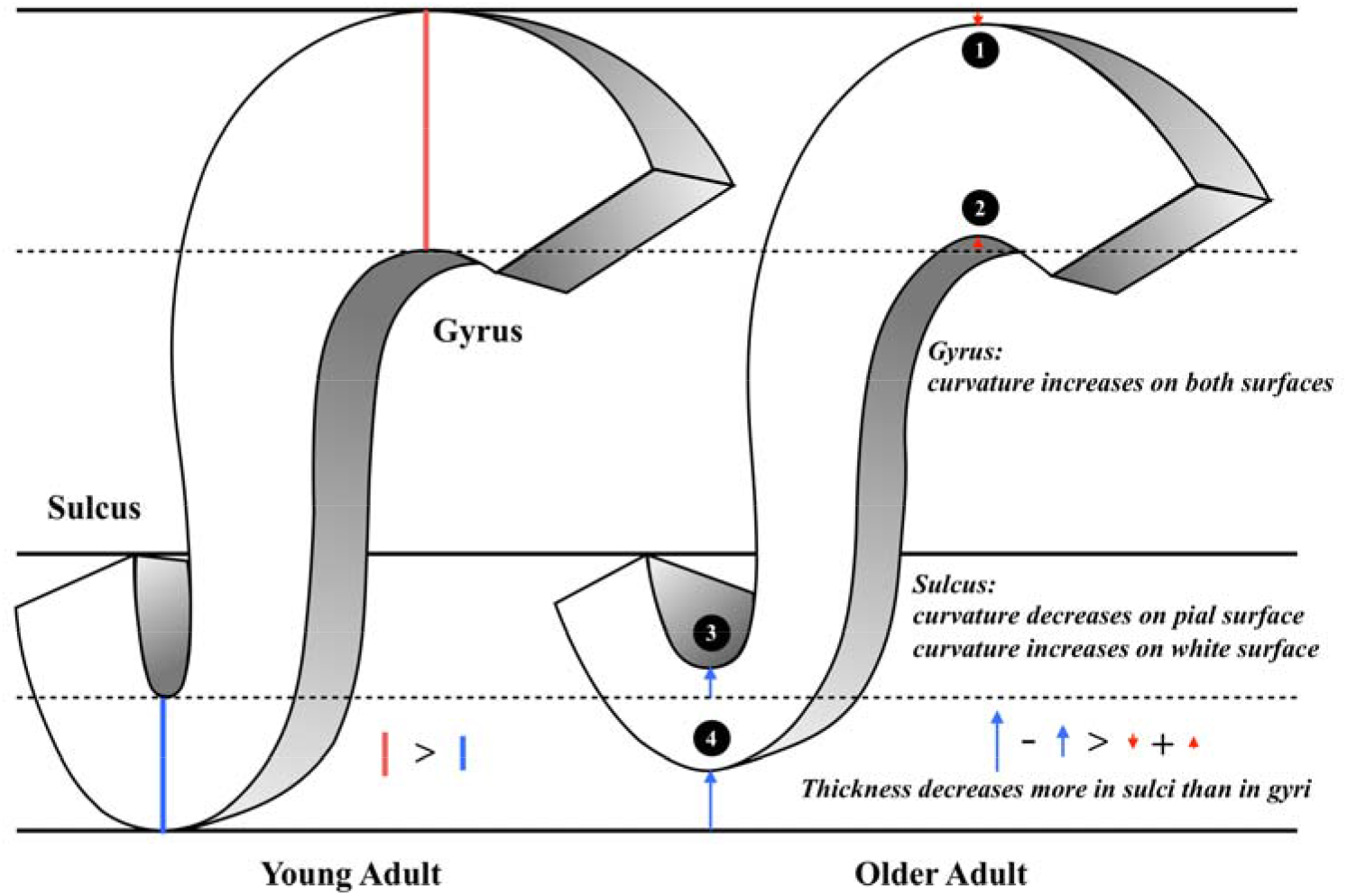
The illustration of the putative morphological model of aging. This figure concludes our main findings as the following: (1) Gyral intrinsic curvature on the pial surface slightly increases with age. (2) Gyral intrinsic curvature on the white matter surface increases with age. (3) Sulcal intrinsic curvature on the pial surface decreases with age; sulcal width increases and sulcal depth decreases with age (4) Sulcal intrinsic curvature on the white matter surface increases with age. (2,4) On the white matter surface, gyral intrinsic curvature increases more intensely with age than sulcal curvature. (1,2,3,4) Sulcal thickness declines more than gyral thickness (see the length of the red and blue arrows).

## Discussion

This study aims to reveal the pattern of the gyral and sulcal changes on the pial and white matter surfaces during the normal aging process using a relatively large cohort dataset. We investigated the whole-brain vertex-wise pattern of the structural changes during the aging process, in which the white matter and pial surfaces showed a different association of gyrification with age. The gyri/sulci ratio was used to highlight the difference of cortical thinning and curvature changes between gyri and sulci spanning across the lifespan. Current findings suggest that, instead of gyri, the changes of sulcal thickness and curvature contribute more during the normal aging process. Finally, we also concluded a putative model of aging in this study based on previous evidence for the fundamental shape of the cortex that accompanied by the current results, which provides a better understanding of the cortical structure degeneration across the lifespan.

As the advance of age, we observed that the curvature decreased on the sulcal-pial surface and increased on the sulcal-white matter surface (Fig. 2). Previous studies have found that sulcal width increases while sulcal depth decreases with age (Kochunov et al., 2005; Liu et al., 2013). This finding partially supports the fact that the sulcal-pial surface might flatten and move outward instead of shrinking toward the white matter surface, which is mainly caused by the steady production of CSF from the choroid plexus that inflates the cerebrum during the loss of brain parenchyma (Matsumae et al., 1996; Miller et al., 1987; Scahill et al., 2003). Moreover, the increased curvature of the WM surface is associated with imbalanced WM-GM shrinkage (Deppe et al., 2014). The specific changes of sulcal morphology in WM-GM surface can be linked with the loss of short association fibers (U-fiber) underneath sulcus that contribute to the impaired local clustering of the brain connections (Gao et al., 2014; Toro and Burnod, 2005; Van Essen et al., 2018). Third, by analyzing the gyri/sulci ratio, we found that the degree of sulcal thickness thinning was larger than that of gyral thickness during the normal aging process. These findings implied that the changes of sulcal morphology are more prominent than gyral regions during aging, which resulted in the variation in the following ways: 1) For sulcal morphology, while the sulcal pial surface moved outward and flatten, the GM-WM surface became cusped and the thickness of sulci decreases. 2) For gyral regions, the curvature on both surfaces became cusped, with mildly decreased gyral thickness. All inferences and current evidence were integrated and graphed in the putative model of aging (Fig. 4).

One of the noticeable findings in the current study is that gyri and sulci altered differently during the normal aging process. Gyral crowns were reported having specialized and enhanced connections and organization between cortices (Brodmann, 1909; Welker, 1990). Although common functional regions were mostly defined in the gyral region and its adjacent sulci, our findings suggested that the sulci itself greatly altered and may be responsible for the decrease of functional segregation during aging. Previous studies using diffusion-weighted imaging have indicated that gyral regions show denser white matter fibers than sulcal regions (Chen et al., 2013; Nie et al., 2012). Deng et al. (2014) further supported these findings that gyri are functional connection centers, while sulci are likely to be local functional units connecting neighboring gyri through inter-column cortico-cortical fibers. Moreover, Gao et al. (2014) found association between the disintegrity in short-range fibers and lower cognitive efficiency on prospective memory, and the loss of the clustering coefficient has also been found to correlate with inferior intelligence quotient (IQ) (Li et al., 2009). Taking together, we suggest that morphological degeneration in sulcal regions could be more vulnerable to the effects of aging. The loss of local interconnectivity in the brain might be more pronounced in the normal aging process. In current study, we found that MMSE is positively correlated with sulcal cortical thickness and gyral intrinsic curvature on the white matter surface. Therefore, the sulcal thickness declines suggest that part of the short-ranged connectivity changes and may influence intra-cortical brain function (Cullen et al., 2010; Elston, 2003; Schuz and Palm, 1989; Wagstyl et al., 2015). However, targeting the cognitive correlates of sulcal degeneration and its impact to brain topology should be investigated in future studies.

We also found different trends of curvature changing on the WM-GM boundary and the pial surface during aging. Based on the retrogenesis mechanism, the differential proliferation hypothesis (Richman et al., 1975) may support differential degeneration process of gyrification of the pial and white matter surfaces during aging. Changes in myelin and synapses in the cortex could be the reason for the morphology development and degeneration, and thereby resulting in cognitive decline (Bartzokis, 2004; Fjell and Walhovd, 2010; Masliah et al., 2006; Whitaker et al., 2016). Nonetheless, a direct link between cortical myelination and gyrification still needs to be found. Although several hypotheses of the mechanisms underlying gyrification are currently being debated by neuroscientists, some have suggested that gyrification is shaped or influenced by multiple mechanisms (Ronan and Fletcher, 2015). The alterations in aging-related gyrification could reflect the underlying cortical connections and the functions of our brain by either pruning or degenerative processes (Jockwitz et al., 2017; Ronan et al., 2011). The vulnerability of sulcal cortical thinning that was discovered in the current study could shed light on the structure-function relationship in the human brain.

The aging model of cerebral cortex we proposed in this study is a global effect spanning across most of the brain regions. However, while most of the regions showing greater gyri/sulci ratio, frontal lobe and cingulate showed similar degree of a decrease between gyral and sulcal thickness (Table 3). Our findings implied that the gyral and sulcal thickness in the frontal and cingulate regions decreased to the same extent during the normal aging process. Several studies have found accelerated regional gray matter volume decline in the frontal lobe compared with other lobes (Resnick et al., 2003; Tisserand et al., 2002) and have demonstrated decreases in cortical volume specifically in the frontal and cingulate regions. These findings consistently suggest that the frontal and cingulate cortices may be the key regions in the brain aging process. Another study reported a higher increase of sulcal width in the superior frontal sulci and a lower correlation between age and decreased sulcal depth in the inferior and orbitofrontal sulci (Kochunov et al., 2005). This report might support our findings that both gyral and sulcal thickness decreased to the same degree so that sulcal width increased significantly and sulcal depth decreased because of the high atrophy in gyri. Moreover, the trend for sulcal gyrification changes on the white matter surface in the occipital lobe did not increase as we found in the whole-brain examination. Small variations exist in different brain regions because the development of the lobes and their functions are diverse.

In this study, we characterized the effects of age on different structural progressions in a large sample of healthy adults. However, this study had several limitations. First, the causality of cortical thickness and intrinsic curvature affecting brain function was not investigated. Due to the nature of the cross-sectional design, we were unable to avoid cohort effect or indicate which structure degenerated earlier or had a higher impact on the brain function. The structure-function relationships among gyri, sulci, and the pial and white matter surfaces and how they impact each other still need to be examined. In this case, the envisioned model of the degeneration of the cerebral cortex may need to be investigated with longitudinal data. Second, to generalize the degenerative process, the current model focused on the trends for comprehensive changes in the cortex. However, cortical variations during aging, including changes in volume and thickness, have been found to decline regionally (Thambisetty et al., 2010; Westlye et al., 2010). Therefore, although we posited general aging-related alterations in the gyri and sulci of the cortex in the model, regional variations still need to be specified. Lastly, participants with mild or severe cognitive functions were excluded in this study, and only general cognition assessments were conducted for current participants. Thus, the generalizability of current findings is limited to healthy population, moreover, its cognitive implications should be further examined with detailed cognitions such as verbal memory, visual executive in future studies.

## Conclusion

This study illustrates a cortical degeneration model from the perspective of brain morphology which provides an overview for the brain aging process using multiple structural measurements,. We found systematical and nonuniform cortical thinning during normal aging, that the overall degree of sulcal degeneration is greater than gyri in terms of thickness and gyrification. These degeneration mechanisms might relate to pruning, life-long reshaping and neurodegenerative processes, associating with differential brain functional degeneration and the underlying neuronal tension. We suggest that the cortical features of gyri, sulci, the pial and white matter surfaces should be considered independently in future studies, which could be associated with segregation and integration alterations in brain connectome during aging.

## Declaration

### Ethical Approval

All procedures performed in the studies involving human participants were in accordance with the Declaration of Helsinki and approved by the Institutional Review Board of Taipei Veterans General Hospital.

