## Supplementary for "Differential patterns of gyral and sulcal morphological changes during normal aging process"

Hsin-Yu Lin^1,2†^, Chu-Chung Huang^1,3*^, Kun-Hsien Chou^1,4^, Albert C. Yang^5,6^, Chun-Yi Zac Lo^3^, Shih-Jen Tsai^5,7^, Ching-Po Lin^1,3,4*
1^Institute of Neuroscience, National Yang-Ming University, 11221 Taipei, Taiwan

^2^Centre for Research and Development in Learning, Nanyang Technological University, 639798, Singapore

^3^Institute of Cognitive Neuroscience, School of Psychology and Cognitive Science, East China Normal University, Shanghai, China

^4^Brain Research Center, National Yang-Ming University, 11221 Taipei, Taiwan

^5^Department of Psychiatry, Taipei Veterans General Hospital, 11221 Taipei, Taiwan

^6^Division of Interdisciplinary Medicine and Biotechnology, Beth Israel Deaconess Medical Center / Harvard Medical School, Boston, MA 002215, USA

^7^School of Medicine, National Yang-Ming University, 11221 Taipei, Taiwan

^†^ These authors contributed equally to this work.

***Corresponding authors:**

Dr. Chu-Chung Huang

Institute of Cognitive Neuroscience, School of Psychology and Cognitive Science, East China Normal University, Shanghai, China

Dr. Ching-Po Lin

Institute of Neuroscience, National Yang-Ming University, 155, Li-Nong St. 11221, Taipei, Taiwan

**Supplemental Materials**

**Linear regression results**

Result of age correlated with cortical thickness showed in Figure S1. Both gyral and sulcal regions negatively correlated with age. The adjusted R-square values were 0.400/0.363 and 0.596/0.559 for left/right hemisphere respectively, which were both moderately correlated. Gyri/Sulci thickness ratio increased while the age increased and the adjusted R-square were 0.320/0.262.

The aging trends of gyral and sulcal intrinsic curvature on pial surface were opposite. In gyral region, intrinsic curvature was slightly increasing, hardly altered with age (R^2^=0.054/0.051) while it was declining in sulcal regions (R^2^=0.474/0.442). Therefore, gyri/sulci ratio increased by age and had R-square values 0.375/0.334. On the gray-white matter boundary of cortical cortex, both gyral and sulcal intrinsic curvature increased with age. Their R-square values were 0.467/0.452 in gyral region and 0.316/0.334 in sulcal region. The gyri/sulci ratio remained stable and unchanged during lifespan. All permutation tests were significant (p < 0.05).

**Supplemental Figures**

**Fig. S1** Pearson correlation between age and GMV / WMV / CSF / TIV. *p<0.05

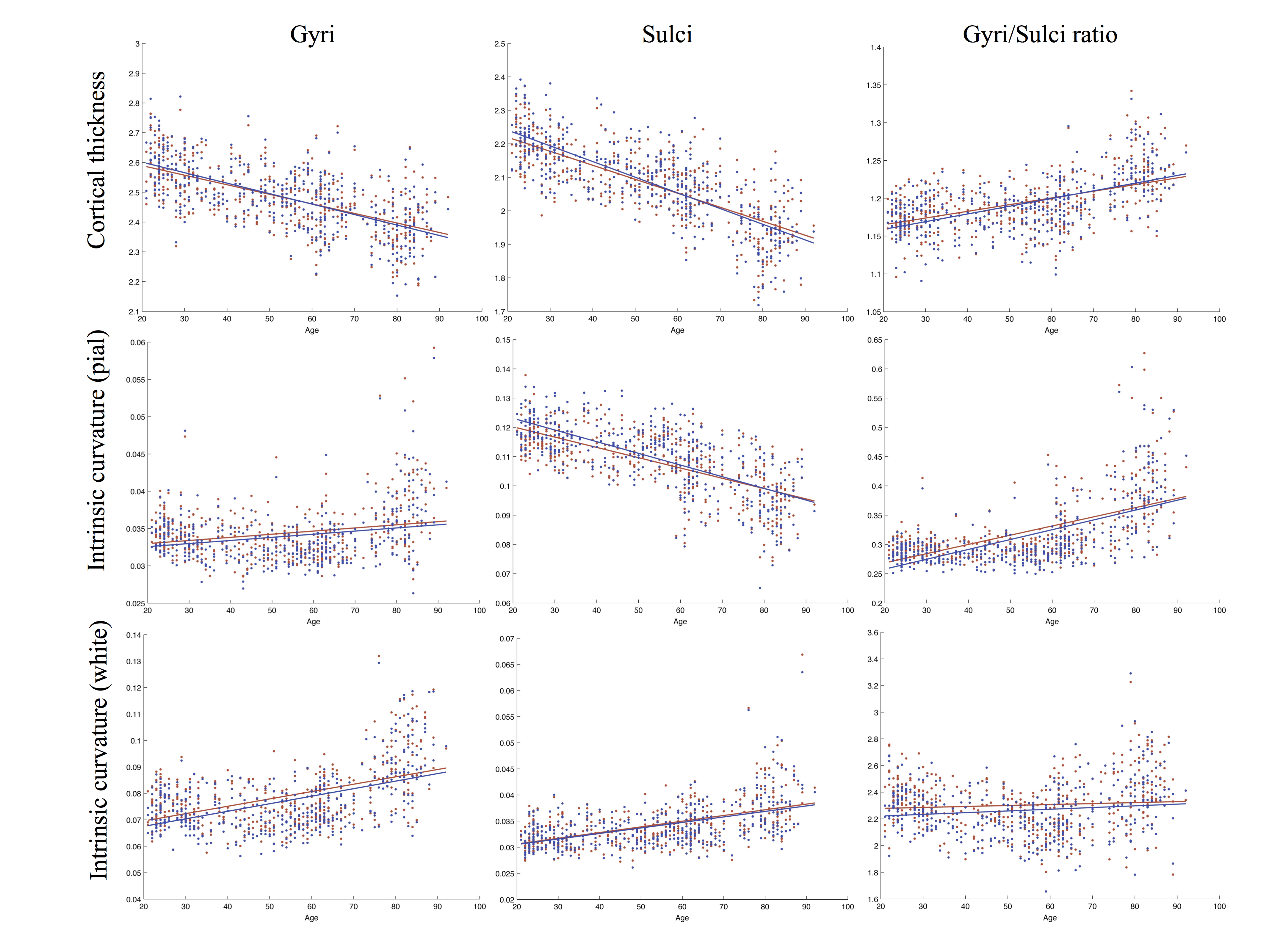

**Fig. S2** Linear regression results of age with gyri, sulci and gyri/sulci ratio of cortical thickness and intrinsic curvature on pial/white surfaces. The red and blue color represent the right and left hemisphere respectively. The lines refer to the fitting curve for the age and measurements of thickness, and the dots indicate the distribution of that data for the subjects.

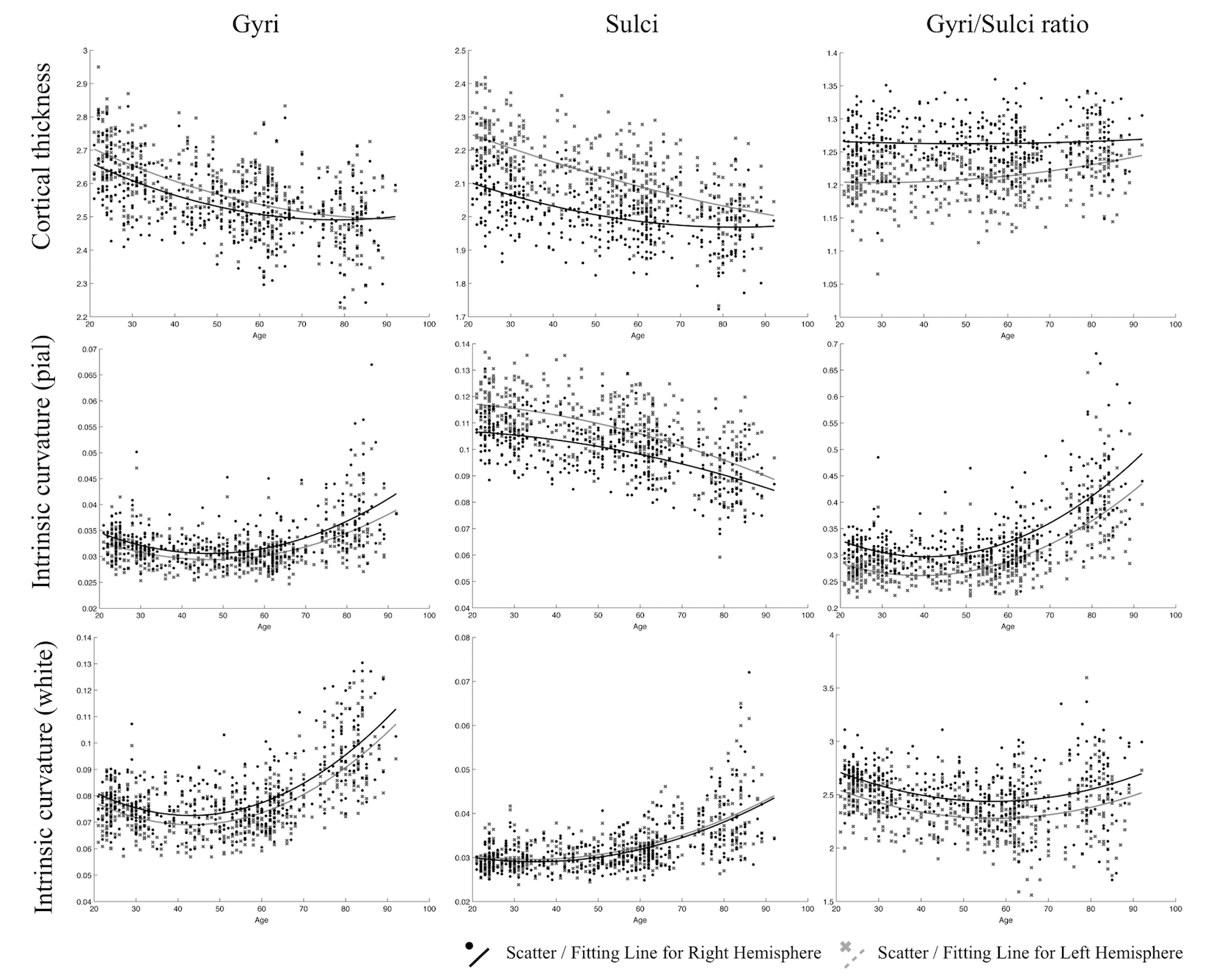

**Fig. S3** Quadratic regression results in frontal lobe of age with gyri, sulci and gyri/sulci ratio of cortical thickness and intrinsic curvature on pial/white surfaces. The lines refer to the fitting curve for the age and measurements, and the dots indicate the distribution of that data for the subjects.

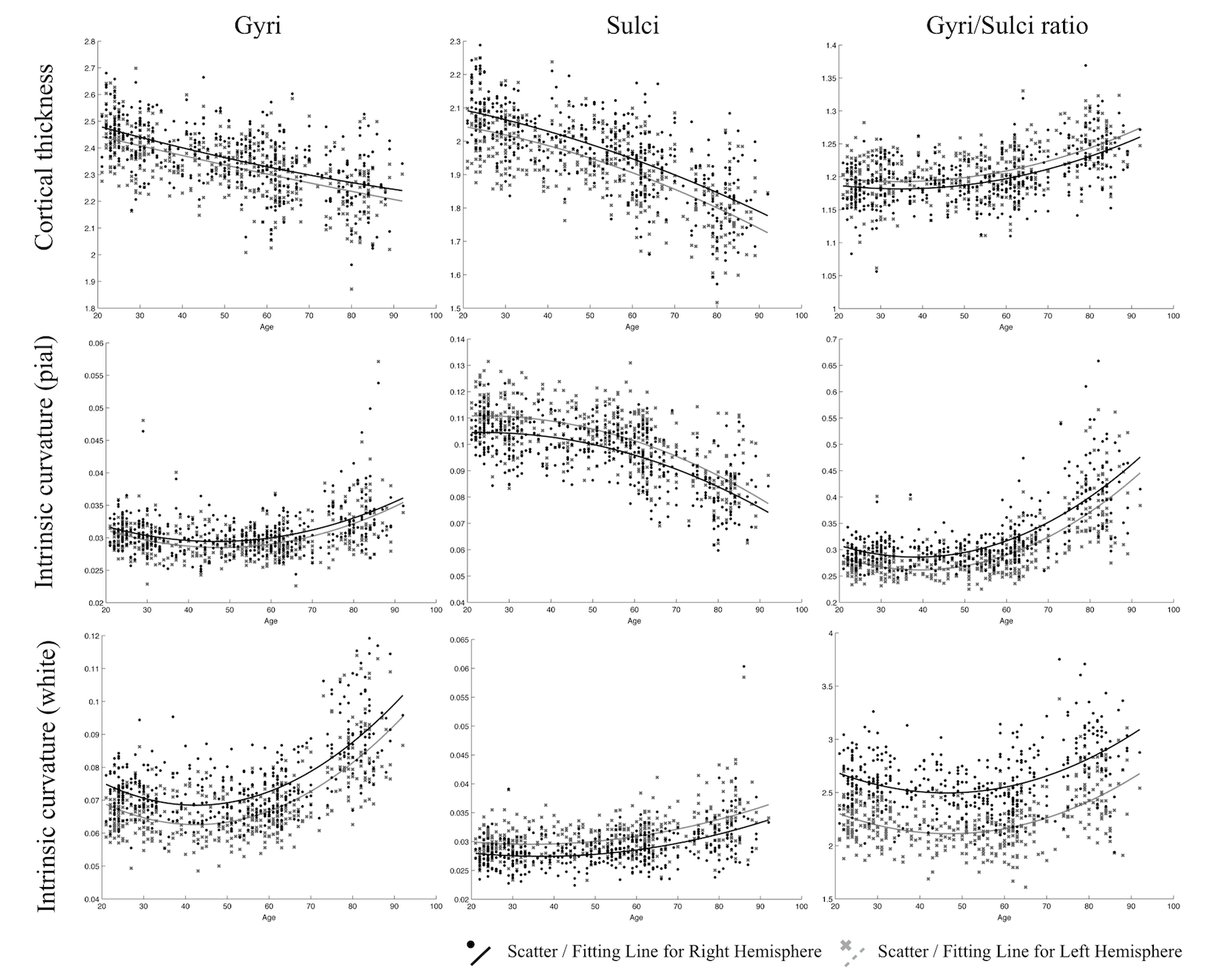

**Fig. S4** Quadratic regression results in parietal lobe of age with gyri, sulci and gyri/sulci ratio of cortical thickness and intrinsic curvature on pial/white surfaces. The lines refer to the fitting curve for the age and measurements, and the dots indicate the distribution of that data for the subjects.

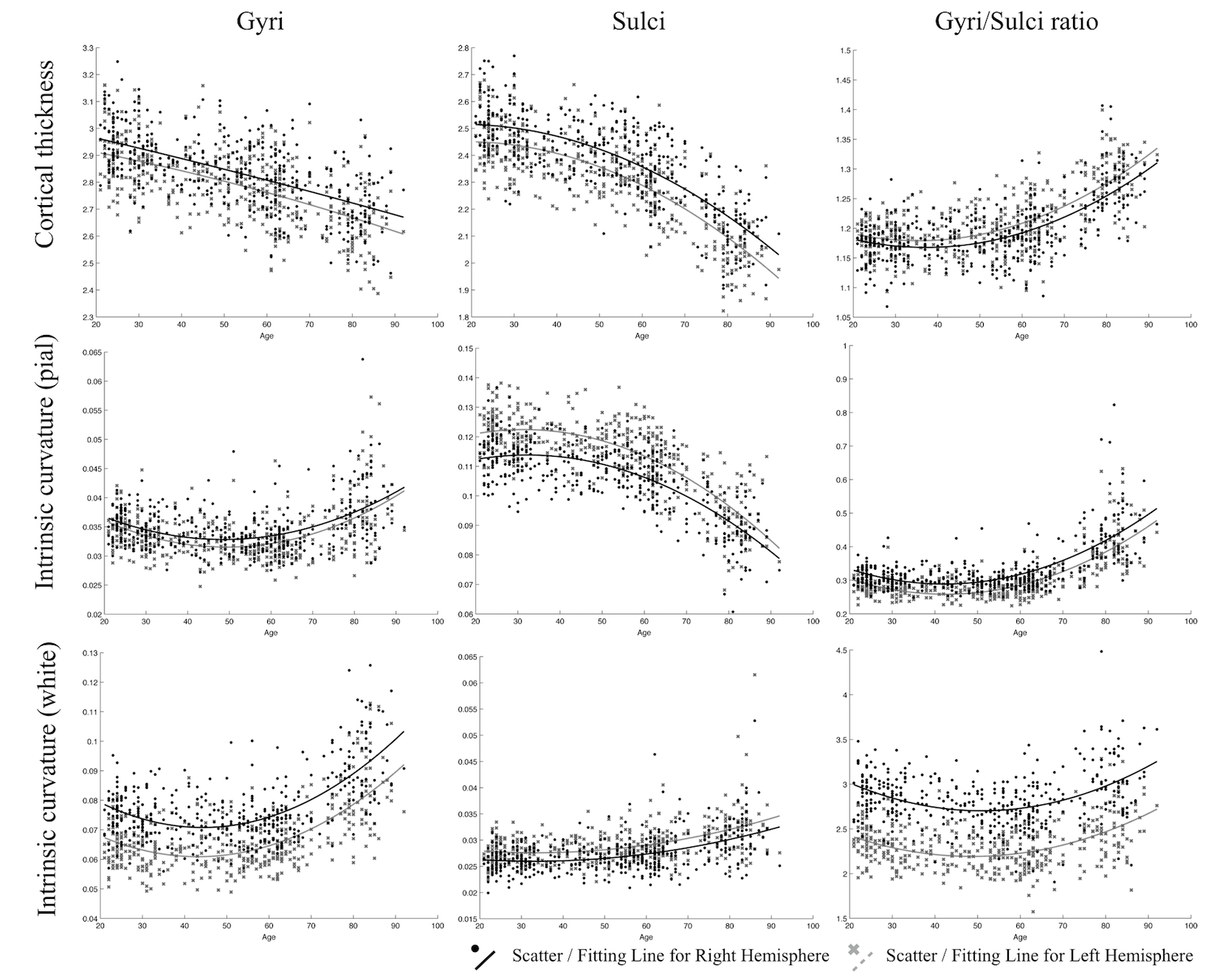

**Fig. S5** Quadratic regression results in temporal lobe of age with gyri, sulci and gyri/sulci ratio of cortical thickness and intrinsic curvature on pial/white surfaces. The lines refer to the fitting curve for the age and measurements, and the dots indicate the distribution of that data for the subjects.

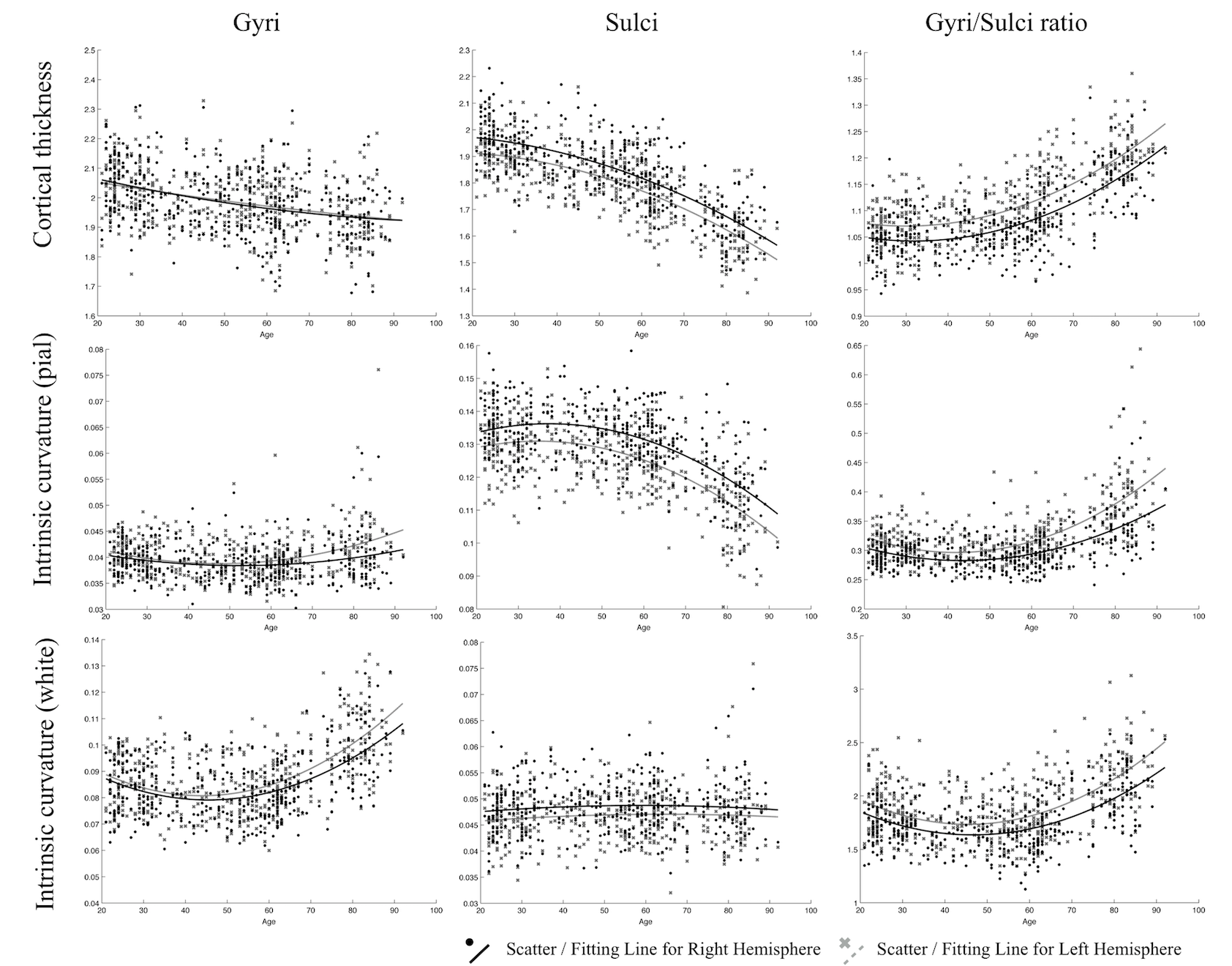

**Fig. S6** Quadratic regression results in occipital lobe of age with gyri, sulci and gyri/sulci ratio of cortical thickness and intrinsic curvature on pial/white surfaces. The lines refer to the fitting curve for the age and measurements, and the dots indicate the distribution of that data for the subjects.

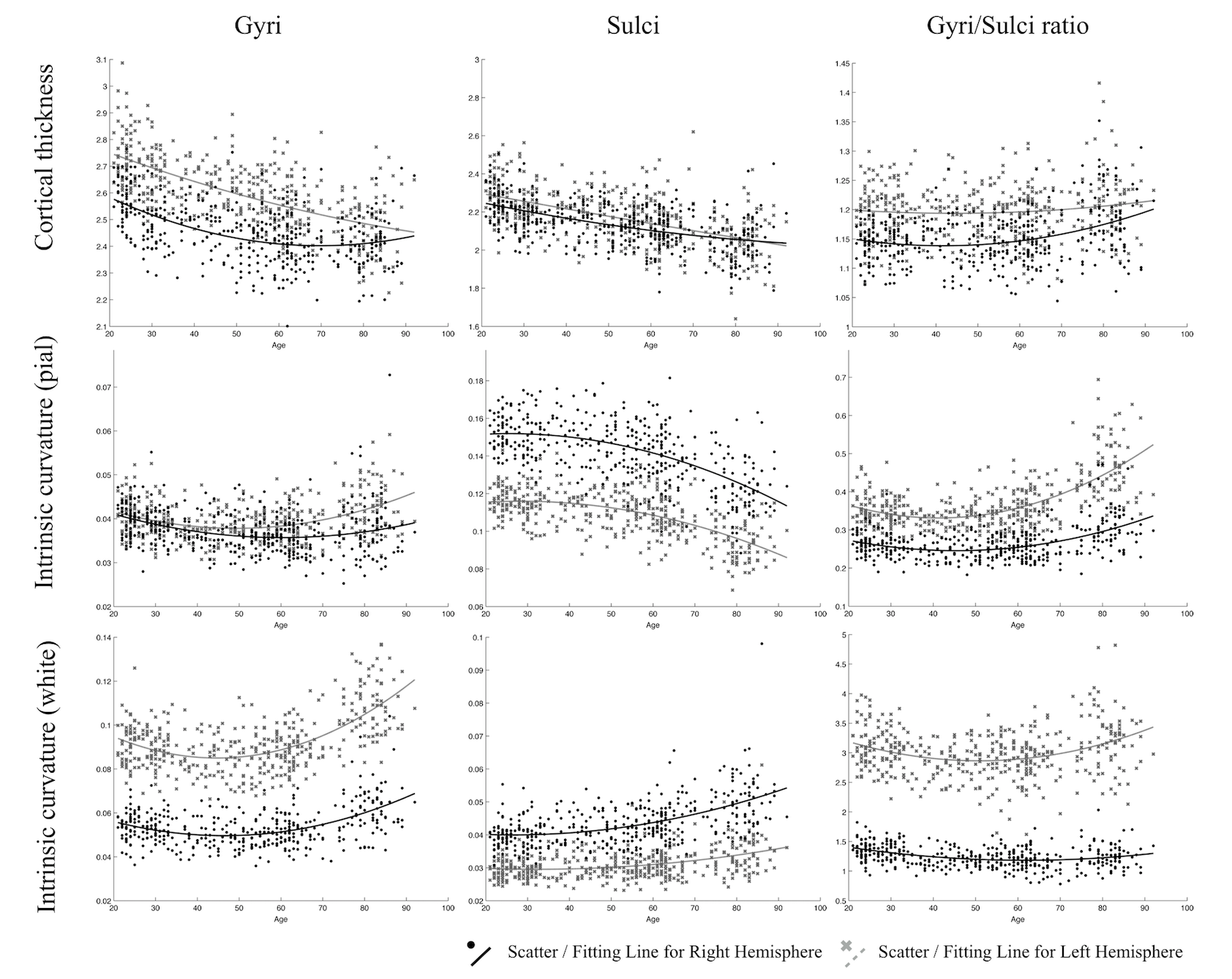

**Fig. S7** Quadratic regression results in cingulate of age with gyri, sulci and gyri/sulci ratio of cortical thickness and intrinsic curvature on pial/white surfaces. The lines refer to the fitting curve for the age and measurements, and the dots indicate the distribution of that data for the subjects.

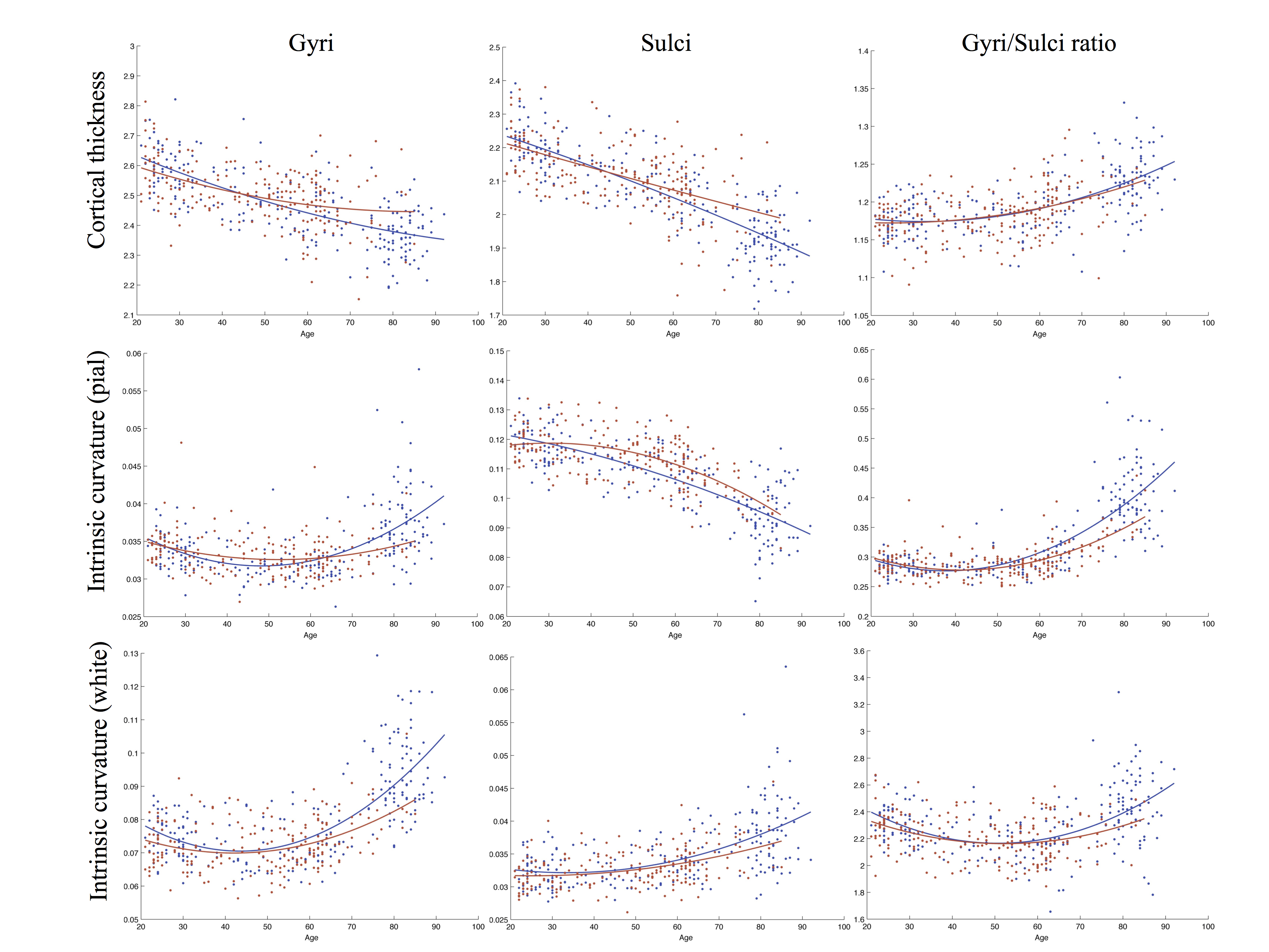

**Fig. S8** Quadratic regression results of age with gyri, sulci and gyri/sulci ratio of cortical thickness and intrinsic curvature on pial/white surfaces grouped by sex on left hemisphere. The red and blue color stood for female and male respectively. The lines refer to the fitting curve for the age and measurements, and the dots indicate the distribution of that data for the subjects.

**Supplemental Tables**

**Table S1** Goodness of fit – Cortical thickness

| Model | RMSE (lh/rh) | AIC value (lh/rh) |
| --- | --- | --- |
| Gyri | | |
| Linear | 0.08809 / 0.08693 | -885.2 / -896.8* |
| Quadratic | 0.08812 / 0.08675* | -883.9 / -897.6 |
| Cubic | 0.08819 / 0.08683 | -882.2 / -895.8 |
| Sulci | | |
| Linear | 0.07884 / 0.07592 | -982.5 / -1015.7 |
| Quadratic | 0.07799 / 0.07547* | -991.2 / -1019.9* |
| Cubic | 0.07805 / 0.07542 | -989.4 / -1019.5 |
| Gyri/Sulci ratio | | |
| Linear | 0.03034 / 0.03040 | -1821.0 / -1819.3 |
| Quadratic | 0.02881 / 0.02895* | -1865.4 / -1861.3* |
| Cubic | 0.02885 / 0.02893 | -1863.4 / -1860.8 |

*RMSE,* Root Mean Square Error: lower is better

*AIC* Akaike Information Criterion: lower is better

* better fitting properties

**Table S2** Goodness of fit - Intrinsic Curvature on pial surface

| Model | RMSE (lh/rh) | AIC value (lh/rh) |
| --- | --- | --- |
| Gyri | | |
| Linear | 0.003519 / 0.003628 | -3712.5 / -3685.8 |
| Quadratic | 0.003139 / 0.003278 | -3811.7 / -3773.8 |
| Cubic | 0.003124 / 0.003262* | -3815.0 / -3777.1* |
| Sulci | | |
| Linear | 0.008598 / 0.008090 | -2928.1 / -2981.6 |
| Quadratic | 0.008424 / 0.007926* | -2945.1 / -2998.6* |
| Cubic | 0.008433 / 0.007934 | -2943.2 / -2996.7 |
| Gyri/Sulci ratio | | |
| Linear | 0.04443 / 0.04544 | -1486.2 / -1466.4 |
| Quadratic | 0.03836 / 0.03992 | -1614.1 / -1579.2 |
| Cubic | 0.03826 / 0.03983* | -1615.5 / -1580.2* |

*RMSE,* Root Mean Square Error: lower is better

*AIC* Akaike Information Criterion: lower is better

* better fitting properties

**Table S3** Goodness of fit - Intrinsic Curvature on white surface

| Model | RMSE (lh/rh) | AIC value (lh/rh) |
| --- | --- | --- |
| Gyri | | |
| Linear | 0.010130 / 0.010220 | -2784.5 / -2776.4 |
| Quadratic | 0.008741 / 0.008898 | -2912.6 / -2897.0 |
| Cubic | 0.008712 / 0.008868* | -2914.6 / -2899.0* |
| Sulci | | |
| Linear | 0.003658 / 0.003715 | -3678.5 / -3664.9 |
| Quadratic | 0.003513 / 0.003574 | -3713.0 / -3697.9 |
| Cubic | 0.003497 / 0.003563* | -3716.1 / -3699.7* |
| Gyri/Sulci ratio | | |
| Linear | 0.2019 / 0.2015 | -157.0 / -158.6 |
| Quadratic | 0.1849 / 0.1858* | -233.0 / -229.1* |
| Cubic | 0.1851 / 0.1860 | -231.2 / -227.1 |

*RMSE,* Root Mean Square Error: lower is better

*AIC* Akaike Information Criterion: lower is better

* better fitting properties

**Table S4** Results of linear regression analyses

| Age correlation | Cortical Thickness  (lh/rh) |  | Intrinsic Curvature (lh/rh) | | |
| --- | --- | --- | --- | --- | --- |
|  |  |  | Pial surface |  | White surface |
| Gyri | **–** (0.400/0.363) |  | + (0.054/0.051) |  | + (0.467/0.452) |
| Sulci | **–** (0.596/0.559) |  | **–** (0.474/0.442) |  | + (0.316/0.334) |
| Gyri/Sulci ratio | + (0.320/0.262) |  | + (0.375/0.334) |  | + (0.014/0.003) |

The variables are adjusted r-square; + represents positive correlation, **–** represents negative correlation

**Table S5** Average cortical thickness and intrinsic curvature

|  | Cortical Thickness  (lh/rh) |  | Intrinsic Curvature (lh/rh) | | |
| --- | --- | --- | --- | --- | --- |
|  |  |  | Pial surface |  | White surface |
| Gyri | 2.483/2.485 |  | 0.034/0.034 |  | 0.079/0.079 |
| Sulci | 2.082/2.086 |  | 1.109/0.110 |  | 0.034/0.034 |
| Gyri/Sulci ratio | 1.194/1.193 |  | 0.321/0.314 |  | 2.303/2.263 |

The variables are the average values of measurements
